# Diversity of biocrust-forming cyanobacteria in a semiarid gypsiferous site from Central Spain

**DOI:** 10.1101/136390

**Authors:** Concha Cano-Díaz, Pilar Mateo, M. Ángeles Muñoz, Fernando T. Maestre

**Affiliations:** Departamento de Biología, Facultad de Ciencias, Universidad Autónoma de Madrid, Madrid, Spain; Departamento de Biología y Geología, Física y Química Inorgánica, Escuela Superior de Ciencias Experimentales y Tecnología, Universidad Rey Juan Carlos, Calle Tulipán s/n, Móstoles 28933, Spain

**Author notes:** E-mail addresses.

**Keywords:** Soil cyanobacteria, Biocrust, Biological soil crust, *Microcoleus*, Cyanobacterial diversity, 16S rRNA

## Abstract

Cyanobacteria are an important constituent of biocrusts, communities dominated by lichens, mosses and associated microorganisms, which are prevalent in drylands worldwide and that largely determine their functioning. Despite their importance, there are large gaps in our knowledge of cyanobacteria associated to biocrusts, particularly in areas such as the Mediterranean Basin. We studied the diversity of these cyanobacteria in a gypsiferous grassland from Central Spain using both morphological identification after cultivation and genetic analyses with the 16S rRNA gene. Eight different morphotypes were observed, most of them corresponding to filamentous and one of them to unicellular cyanobacteria. We found cyanobacterial genera typical of biocrust communities, such as *Microcoleus*, *Schizothrix* or *Tolypothrix*, and N-fixing cyanobacteria as *Scytonema* or *Nostoc*. Genetic information allowed us to identify cultures belonging to recently described genera such as *Roholtiella*, *Nodosilinea* and *Mojavia*. In this study we describe two new phylotypes of *Microcoleus* and *Scytonema*, which are key genera contributing to ecosystem functioning in biocrust-dominated ecosystems worldwide.

## 1. Introduction

Cyanobacteria are a cosmopolitan and diverse group of microorganisms that are a key component of biocrusts, soil surface communities also formed by lichens, mosses, liverworts and other microorganisms that are a prevalent biotic feature of drylands worldwide (Büdel et al., 2016). Cyanobacteria are present in virtually all biocrust communities due to their capacity to adapt to a wide range of ecological conditions (Tamaru et al., 2005). Early successional biocrusts are dominated by filamentous cyanobacteria, which are the first colonizers of bare ground areas in drylands (Garcia-Pichel and Wojciechowski, 2009). These organisms segregate an extracellular polysaccharide (EPS) matrix that promotes soil stabilization and enhances microhabitat conditions for colonization of other cyanobacteria and the rest of biocrust constituents (Mager and Thomas, 2011). Heterocyst-forming cyanobacteria are important contributors to carbon and nitrogen fixation in oligotrophic ecosystems such as drylands (Belnap, 2002), and it has been estimated that together with the rest of components of cryptogamic covers they contribute to the fixation of nearly half of the total amount of biologically fixed nitrogen worldwide (Elbert et al., 2012). Given the important ecological roles they play, and the strong links between species composition and diversity and ecosystem functioning in biocrust communities (Bowker et al., 2013; Maestre et al., 2012; Yeager et al., 2012), understanding the composition and diversity of cyanobacterial communities can provide valuable information to assess ecosystem functioning and development in biocrust-dominated landscapes.

Cyanobacterial communities within biocrusts are a common target of interest for researchers around the world. Detailed studies of cyanobacterial species composition and distribution have been carried out in North America, Asia, Africa, and Europe (e.g. Garcia-Pichel et al. 2013, Dojani et al. 2014, Hagemann et al. 2014, Kumar and Adhikary 2015, Williams et al. 2016). However, few studies so far have analysed the cyanobacteria associated to biocrusts in gypsum habitats (Garcia-Pichel et al., 2001; Steven et al., 2013), despite they are hotspots of botanical diversity (Escudero et al., 2014) and harbour very conspicous biocrust communities dominated by lichens (Castillo-Monroy et al., 2010; Martínez et al., 2006). With the aim to advance our knowledge about cyanobacterial communities associated to biocrusts in gypsum soils, we studied them in a gypsiferous semiarid site of central Spain. For doing so we used the combination of molecular and morphological information, as this improves the quality and quantity of sequences in molecular databases, which is a major concern in the study of cyanobacterial diversity nowadays (Thomazeau et al., 2010; Weber et al., 2016).

## 2. Materials and methods

### 2.1 Field site

This study was carried out in the Aranjuez Experimental Station, located in central Spain (40°01'55.7"N - 3°32'48.3"W and 590 m above sea level). Climate is semiarid, with an intense drought in summer from June to September. Mean annual temperature is 15°C and annual precipitation is 349 mm. Soil is rich in gypsum (*Gypsiric Leptosoil*; IUSS Working Group WRB 2014; see Castillo-Monroy et al. 2010 for a physico-chemical characterization). Vegetation is characterized by herbaceous plants like *Macrochloa tenacissima* and shrubs like *Retama sphaerocarpa* and *Helianthemum squamatum*. Soil among vegetation is covered by a well-developed biocrust community dominated by the squamous lichens *Diplochistes diacapsis*, *Squamarina lentigera* and *Psora decipiens*; with patches of acrocarpous mosses *Pleurochaete squarrosa* and *Tortula revolvens* (see Maestre *et al.* 2013 for a full list of lichens and mosses found in the site).

### 2.2 Soil collection and morphological characterization of cyanobacteria

We randomly selected eight plots in areas with a well-developed biocrust community in July 2013. At each plot, we took five samples (0-1 cm depth), which were pooled and taken to the lab. Lichens and mosses were removed and soil was sieved through a 2 mm sieve and kept dry in darkness conditions.

Cyanobacterial strains were isolated using a modification of the procedure described in Loza et al. (2013). Aliquots of ∼1 g of soil were mixed with 1.5 ml of cyanobacterial culture media and distributed uniformly over different solid media (1.5% agar concentration). We used four common culture media for cyanobacteria: BG11, BG11_0_ (Rippka et al., 1979), modified CHU 10, and modified CHU 10 without addition of N (Gómez et al., 2009). These media allowed the growth of cyanobacteria by providing a range of nutrient richness and absence or presence of N, which is important to isolate both, N-fixing cyanobacteria as well as non-N-fixing cyanobacteria. To avoid fungal contamination, we added cycloheximide (0.1 mg/ml). Cultures were incubated in a growth chamber at constant light and temperature (20-50 µmol photons m^-2^ s^-1^ and 28°C) three to four weeks until colonies grew without overlapping. Cyanobacterial colonies were picked up under a dissecting microscope (Leica, Leica Microsystems, Wetzler, Germany) with pulled capillary pipettes and forceps to isolate colonies, filaments and bundles(Gómez et al., 2009). Cultures were kept in the same medium and conditions both in agar plates and in liquid medium for further growth.

All colonies were characterized morphologically using a dissecting microscope and an Olympus BH2-RFCA photomicroscope (Olympus, Tokio, Japan). Identification and morphological characterization of the cyanobacteria were performed taking into consideration features relating to colony morphology, trichome shape, sheaths and details of cell morphology. Taxonomy was based on Geitler (1932), Anagnostidis and Komárek (1999), Komárek and Anagnostidis (2005) and Komárek (2013).

### 2.3 Genotypic characterization

DNA was extracted with the Ultraclean Microbial DNA Isolation Kit (Mobio, Carlsbad, CA, USA) following the manufacturer's instructions. A prior step was added at the beginning of the procedure as previously described (Loza et al., 2013): samples were homogenized and exposed to three cycles of thermal shock alternating immersion in liquid N and heating to 60°C to break the protector mucopolysaccaride that covers the surface of many cyanobacteria. PCR amplifications were performed by using the bacterial 16S rRNA primers 27F and 1494R (Neilan et al. 1997). The PCR mixture (25µl) contained 2.5 µl Buffer 10X, 1.5 mM MgCl_2_, 50 µM dNTP, 10 pmol of each primer, BSA 1 mg/ml, 5 µl TaqMasterTM PCR Enhancer 5x (Eppendorf, Germany), 0.75 U Ultratools DNA polymerase (Biotools, Spain), miliQ H_2_O and 10 ng DNA. Amplification took place in a termocycler PCR Eppendorf Mastercycler (Eppendorf, Viena) with the reaction conditions described by Gkelis et al. (2005). Success in PCR was checked by agarose gel 1.5% using 1Kb Gene Ruler (MBL Biotools, Spain) and fluorescent DNA stain GelRed^TM^. PCR products were purified with Real Clean Spin Kit (Real, Durviz, Spain) and sequenced at Centro Nacional de Investigaciones Oncológicas (Madrid, Spain). When sequences did not have enough size (<200 pb) or quality, PCR products were cloned into pGEM-T vectors with the pGEM Easy Vector system (Promega, US) according to manufacturer recommendations and sequenced according to vector information and primers. Sequences were obtained for both strands independently.

We compared our sequences with sequences from the National Center for Biotechnology Information (NCBI) database to complement identifications. For phylogenetic analysis, sequences were aligned using ClustalW (Thompson et al., 1994) with the software Bioedit 7.2.5 (Ibis Biosiences, Carlsbad, CA). We obtained the most similar sequences from the NCBI database (*blast.ncbi.nlm.nih.gov*) with BLAST (Altschul et al., 1990) and then performed multiple alignment with all sequences. Phylogenetic trees were generated with MEGA7 software (Tamura et al., 2013) using the Maximum Likelihood, Neighbor Joining (Saitou and Nei, 1987), and Maximum Parsimony methods and the Tajima Nei matrix (Tajima and Nei, 1984) to calculate pairwise distances. Statistical significance was performed with bootstrap test from Felsenstein (1985) for 1000 replicates in the case of Neighbor Joining tree and 500 replicates for Maximum Parsimony and Maximum Likelihood trees.

Cultures were named after the site (Aranjuez, AR-) and were included in the culture collection of the Universidad Autónoma de Madrid (UAM). The nucleotide sequences obtained in this study were uploaded to the Genbank database (accession numbers: MF002044 - MF002062).

## 3. Results

Macroscopic and microscopic evaluation of cultivated cyanobacteria resulted in the identification of eight different cyanobacterial morphotypes and the successful isolation and sequencing of 12 strains. Three main types of morphologies were found: filamentous and heterocyst-forming cyanobacteria (*Tolypothrix-*type*, Nostoc-*type*, Scytonema*), filamentous cyanobacteria without heterocysts (*Microcoleus-*type, *Leptolyngbya-*type, *Schizothrix*) and unicellular cyanobacteria (*Chroococcus*) (Figure 1; Table 1)

**Fig. 1.**
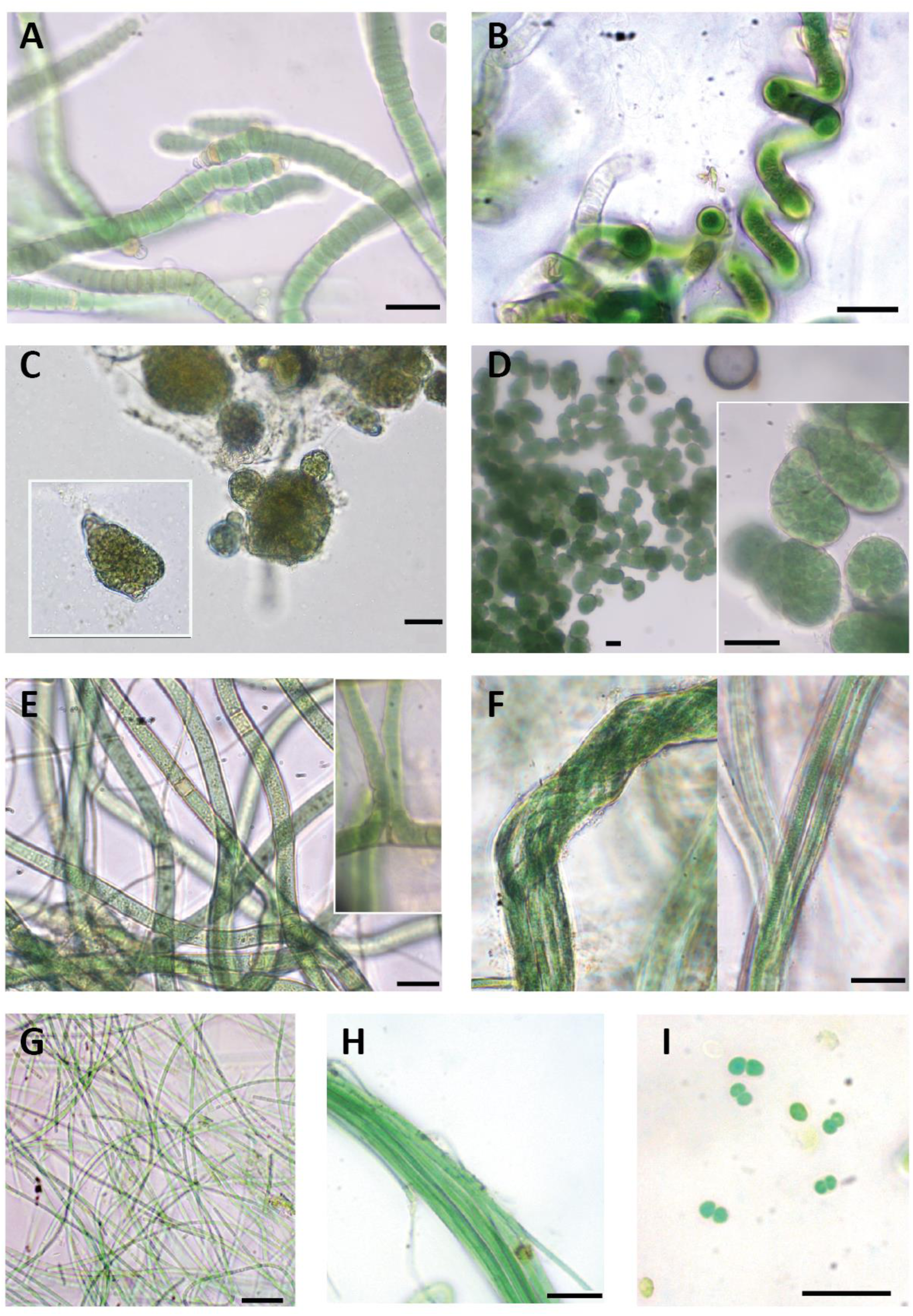
Diversity of cyanobacterial morphotypes found in our study area. (A) *Roholtiella edaphica*. AR2; (B) *Spirirestris* sp; (C) *Nostoc sp.* AR12; (D) *Mojavia* sp. AR1; (E) *Scytonema* sp. In cultivation progress; (F) *Microcoleus* sp. AR10; (G) *Leptolyngbya* sp. AR11,; (H) *Schizothrix* sp.; (I) *Chroococcus* sp. Scale bars equal 20 µm.

**TABLE 1:** Morphological characteristics of the cyanobacterial strains observed in our study. B=breadth (µm); L= cell length (µm)

| Taxon | Culture-Strain | Morphological description | Dimensions ($\mu\text{m}$) | Figure |
| --- | --- | --- | --- | --- |
| *Roholtiella edaphica*. | UAM 802- AR2<br>UAM 803 -AR3<br>UAM 804 -AR4<br>UAM 805 -AR5<br>UAM 806 -AR6 | Heterocystous filamentous. Trichomes into a thin and transparent sheath. False branching occurs. Terminal and apical heterocytes. Rounded to flattened cells, sometimes attenuating its size towards the end of the trichome. | B=6.6-11.3<br>L=3.3-6 | 1 A |
| *Spirirestris* sp. | - | Thick filaments, that turn into spiralized and irregular shapes. Intermediate heterocytes. Rectangular cells, densely packed towards the end of the filament | B= 8-9.3<br>L=3.3-5.3 | 1 B |
| *Nostoc* sp. | UAM 812- AR12 | Heterocystous filamentous. Short chains of cells. Irregular to sphaerical colonies. Surrounded by a thin mucilage. Terminal or lateral heterocytes. Green to yellowish colour. Irregular cells, sometimes sphaerical. | B=5-7<br>L= 5.5-7.5 | 1 C |
| *Mojavia* sp. | UAM 801- AR1 | Heterocystous filamentous. Short chains of cells. Subsphaerical to ellipsoid, irregular colonies surrounded by a firm mucilaginous covering. Terminal or lateral heterocytes. Intensely colored green. Sphaerical to irregular cells, densely aggregated | B=4.3-6.4<br>L= 5.8-6.5 | 1 D |
| *Scytonema* sp. | UAM 807 -AR7<br>UAM 808 -AR8 | Heterocystous filamentous. Thick and cylindrical trichomes, yellow, blue-green and light green. Single and double false branching. Terminal and intermediate heterocysts. Necridia occurs in some filaments. Very thin and transparent sheath. Rectangular cells densely packed towards the end of the filament | B=7.5-9<br>L=3-5 | 1 E |
| *Microcoleus* sp. | UAM 809 -AR9<br>UAM 810 -AR10 | Ensheathed, bundle-forming filaments, surrounded by a very thick and transparent sheath. Sometimes 5-6 thrichomes but also singular filaments. No heterocytes. | B=4.2-5<br>L=1.4-2.1 | 1 F |
| *Nodosilinea* sp. | UAM 811 -AR11 | Single, Long, thin and flexuous filaments. Colorless to Light green color. No heterocytes or akinetes. Cells isodiametrical or longer than wide, cylindrical | B=2.2-3.2<br>L=3.5-5 | 1 G |
| *Schizotrix* sp. | - | Ensheathed, bundle forming thick filaments, containing several trichomes. Sheaths fine and not colored. Intense green and cylindrical trichomes, slightly attenuated at the end. Cylindrical cells, longer than wide. Apical cells are conical. | B=1.7-3.2<br>L=4.5-5.1 | 1 H |
| *Chroococcus* sp. | - | Single cells, aggregated into colonies usually with two cells surrounded by a transparent sheath. Light green | B=2-3.3 | 1 I |

Phylogenetic trees were carried out using our nineteen 16S rRNA sequences together with 37 cyanobacterial sequences from NCBI database. Since the three methods used (Maximum Likelihood, Neighbor Joining and Maximum Parsimony) produced similar clustering only the Maximum Likelihood tree is presented, with an indication of the bootstrap values for all three approaches (Figure 2). Heterocystous cyanobacteria formed a well-supported group separated in four different clusters, and filamentous non-heterocystous strains were included in other two clusters.

**Fig. 2.**
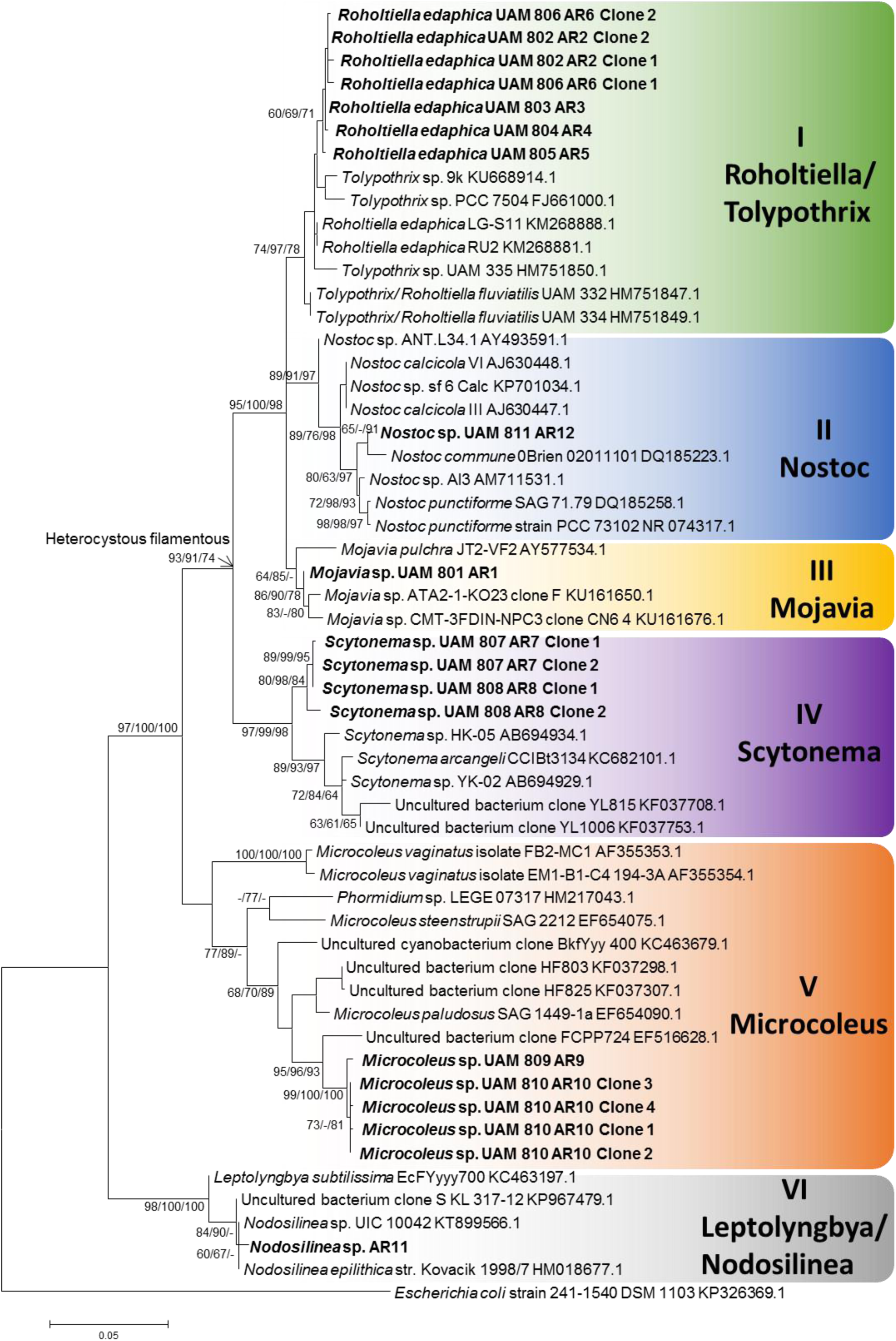
Phylogenetic tree based on 16S rRNA gene sequences obtained by the Maximum Likelihood method (log likelihood -5159.3732). The percentage of trees in which the associated taxa clustered together (Bootstrap) is shown next to the branches (>60% values are reported for Maximum Likelihood, Neigbor Joining and Maximum Parsimony analysis). Sequences from isolated cyanobacteria cultures are in bold. The tree is drawn to scale, with branch lengths measured in the number of substitutions per site. The 0.05 bar indicates substitutions per nucleotidic position.

### Tolypothrix-type

Five isolated strains (AR2 to AR6) were initially identified as *Tolypothrix* sp. They all shared typical characteristics of the genus such as single false branching, as well as terminal and intercalary heterocysts (Figure 1 A). Thrichomes were surrounded by a thin and transparent sheath. In some cases, cells attenuated their size towards the end of the thrichome. Sequences of these isolated strains were included in Cluster I together with sequences of *Tolypothrix* and *Roholtiella* genera ranging from 97.7 to 99.3% sequence similarity (Figure 2). No genetic differences were found between the sequences of our strains (99.8% of similarity) and a 99.3% of similarity with the database sequence from *Roholtiella edaphica*, a recently described Tolypothrichaceae species (Bohunická et al., 2015); therefore these strains can be assigned to this taxon.

Thick cyanobacterial filaments that turn into characteristic *Spirirestris* spiralized forms were observed during the culturing progress but could not be isolated. Intercalary heterocysts also occurred (Figure 1 B). The genus *Spirirestis* shares morphological characters with the genus *Tolypothrix*, principally heterocyst formation, false branching, and presence of the sheath, but most trichomes are tightly spiraled (Flechtner et al., 2002).

### Nostoc-type

Two filamentous and heterocyst-forming isolated strains (AR1 and AR12) were initially identified as belonging to genus *Nostoc* according to traditional morphological criteria (Figure 1 C, 1 D). Isolated strain AR1 had short filaments, with terminal and intercalary heterocysts. Colonies were subsphaerical to irregular, densely aggregated and had an intense green color. A firm mucilaginous sheath covered thickly entwined trichomes (Figure 1 C). Cyanobacterial culture of strain AR12 had a much thinner mucilage and colonies were sphaerical with a green to yellowish color (Figure 1 D). Sequences of these two strains were clearly separated in different clusters of the phylogenetic tree (Cluster II and III) (Figure 2). Cluster II was well supported and included sequences corresponding to *Nostoc* genus with a 98.6-98.8% similarity within the cluster. Isolated strain AR12 showed a similarity of 98.8% with *Nostoc calcicola* from this cluster. Cluster III harbored sequences from *Mojavia* and one *Nostoc*, with 98.3-99.2 % similarity within the cluster. *Mojavia* is a new cyanobacterial genus that shares morphological features with *Nostoc* but has genetic differences in the ITS region (Řeháková et al., 2007). Isolated strain AR1 showed a high similarity (99.2%) with a sequence from the database corresponding to *Mojavia* sp., and therefore culture AR1 can be assigned to this genus.

### Scytonema

Two filamentous and heterocystous cyanobacterial strains (AR7 and AR8) were identified as *Scytonema* sp. They showed the typical characteristics of this taxon such as thick and cylindrical thrichomes, filaments with double and single false branching, and terminal or intercalary heterocysts. Necridia appeared in some filaments and sheath was thin and transparent (Figure 1 E). Cluster IV harbored *Scytonema* sequences (Figure 2) that shared a 97.6-98 % similarity, and had a strong bootstrap support. However, all our sequences from our *Scytonema* strain cultures were located in a sister separated cluster from the other *Scytonema* sequences, with only a maximum of 97% of similarity to the closest relative.

### Microcoleus

Filamentous strains AR9 and AR10 showed morphological features characteristic of *Microcoleus*: typical bundles or dense packages of thrichomes (5-6 thrichomes observed) surrounded by a very thick and transparent sheath, although single-thrichomes in filaments were also observed (Figure 1 F). *Microcoleus* sequences were included in Cluster V (Figure 2) together with other Oscillatoriales such as *Phormidium*, *Microcoleus* as well as uncultured cyanobacteria. *Microcoleus vaginatus* sequences were very different from the rest and formed a well-supported clade, sister to the rest of the group. The nearest relative in the sister branch of our sequences was an uncultured bacterium clone, which exhibited very low similarity (97%). Sequence similarities between representatives of *Microcoleus*, such as *M. paludosus, M. steenstrupii, and M. vaginatus* were only 94.9%, 94.5%, and 91.5%, respectively.

### Leptolyngbya-type

Culture AR11 showed morphological features from the polyphyletic genus *Leptolyngbya*, with single, long, thin and flexuous filaments. No akinetes or heterocytes were found and filaments were light green to transparent (Figure 1 G). Cluster VI grouped sequences from *Leptolyngbya* and *Nodosilinea* that shared high sequence similarity (98.9-99.3%), with high bootstrap support. Strain AR11 exhibited high similarity (99.3%) with *Nodosilinea* sp., a genus that shares morphological characteristics with Leptolyngbya, so culture AR11 can be assigned to this taxon

### Schizothrix

Cyanobacterial cultures with morphological characteristics of *Schizothrix* genus were observed but could not be isolated. Filaments were thick and formed bundles of several trichomes, which were intensely green and slightly attenuated at the end (Figure 1 H). Mucilaginous sheath was fine and not colored.

### Chroococcus

Unicellular cyanobacteria were only observed in a not isolated culture that showed morphological characteristics of *Chroococcus*: single light-green cells, usually aggregated by pairs, surrounded by a transparent sheath (Figure 1 I).

## 4. Discussion

Phylogenetic analysis support the monophyly of heterocystous cyanobacteria (Figure 2), as previously found (Berrendero et al., 2011; Lyra et al., 2001). Percentage of similarity between our sequences and those of the NCBI database were in agreement with phenotypic characteristics, corroborating most of the identifications. Isolated strains AR2-AR6, corresponding to *R. edaphica* were located into a cluster with sequences belonging to *Tolypothrix* genus. *Roholtiella* is a recently described genera (Bohunická et al., 2015) that shares morphological characteristics with other Tolypothricaceae such as *Tolypothrix*, but has genetic differences. The most similar sequence found in the NCBI database is *R. edaphica* LG-S11 from soil samples of Russia (Republic of Bashkortostan). We recorded morphological variability within different cultures (AR2 to AR6) as attenuation of cells towards the end of the thrichome, which were probably due to phenotypic variability frequently observed in cyanobacterial cultures(Berrendero et al., 2011).

Within the *Nostoc*-type, sequence corresponding to strain AR12 was located into an entirely *Nostoc* cluster, so it belongs clearly to this genus. The most similar sequence (98.8%) found in the NCBI database corresponds to a *N. calcicola* from Czech Republic. Strain AR1 was located in a cluster with all sequences from *Mojavia*, a recently described cyanobacterial genus that shares morphological features with *Nostoc* but has genetic differences in the ITS region (Řeháková et al., 2007). Sequence from AR1 is very similar to other *Mojavia* sequences (>99% similarity); however, the only species identified in the cluster, *M. pulchra* JT2-VF2 from dessert crusts of Joshua Tree National Park (USA), differs from our sequence (only 98% similarity).

Our cultures of *Scytonema* also seem to be singular, as they were clustered into the clade of *Scytonema* sequences in our phylogenetic trees, but similarities between our almost identical sequences and the rest from the database are below 98%. The low similarity (97%) of sequences of these strains with the closest relative (*Scytonema* sp. HK-05, isolated from crusts from Japan) suggests a novel biocrust-associated phylotype of *Scytonema*.

Relationships between sequences throughout percentage of similarity and phylogenetic tree have shown the singularity of strains of *Microcoleus* sp. AR9 and AR10. These strains correspond to the most distinct cyanobacteria found in this study. Sequences were located within the *Microcoleus* cluster and had morphological characteristics of this genus, but it seems to be really different from other common *Microcoleus* found in biocrusts such as *M. vaginatus* (only 91% similarity), or *M. steenstupii* (94% similarity). The most similar sequence is not identified or cultured yet and belongs to a grassland soil from northern California (USA) with a 97.5% similarity. For all this reasons we believe we found a new phylotype of *Microcoleus* with no matches in the database.

Isolated strain AR11 sequence falls into the cluster of *Nodosilinea* and *Leptolyngbya*. *Nodosilinea* is another recently described genus (Perkerson et al., 2011) that shows similar morphological characteristics of *Leptolyngbya*, but is genetically and phylogenetically different. Presence of *Leptolyngbya* species in the clade probably is related to their similarity and morphological identifications. In fact, *Leptolyngbya* is a genus in discussion because of its polyphyletic nature.

Studies comparing different parent materials have revealed that gypsum soils have singular cyanobacterial communities (Garcia-Pichel et al., 2001; Steven et al., 2013). In biocrusts from the Colorado Plateau (USA), Garcia-Pichel et al. (2001) found differences in cyanobacterial communities from gypsum soils when compared to sandy, shale and silt soils using denaturalizing gradient gel electrophoresis (DGGE) and microscopy. The common dominant filamentous cyanobacteria *M. vaginatus* appeared in all biocrusts except in those from gypsiferous soils (Garcia-Pichel et al., 2001). In the same area, studies using massive sequencing have found that gypsum soils have a lower abundance of cyanobacteria and diversity than other soils, with very low relative abundances of *M. vaginatus* (1-5%), but being dominant in the rest of the crusts (Steven et al., 2013). In Spain, taxonomic knowledge of biocrust-associated cyanobacteria is poorly developed (Maestre et al., 2011). Using DGGE, Maestre et al. (2006) found 19 different phylotypes in biocrusts from a calcareous site in SE Spain. No heterocystous cyanobacteria were detected, but common crust genera, such as *Leptolyngbya*, *Oscillatoria* or *Phormidium,* and the cosmopolitan *M. steenstupii* were associated to biocrust-forming lichens in this location. We believe that the *Microcoleus* found at our study site could have the same ecological function as other *Microcoleus* as it showed the typical bundles contributing to soil stabilization. The lack of heterocystous cyanobacteria in Maestre et al. (2006) is possibly related to the molecular approach, which can underestimate this part of the cyanobacterial community that commonly appears in cultures (Garcia-Pichel et al., 2001).

Despite culture limitations, as some cyanobacterial genera could not be successfully isolated, and the low number of total different species found, we were able to characterize morphological and phylogenetically twelve isolated strains, belonging to common genera found in soils. Biocrusts at our study site hold a distinctive community of cyanobacteria of principally filamentous forms (heterocystous and non-heterocystous), with representatives of genera typically present in biocrusts from around the world, such as *Microcoleus*, *Scytonema, Nostoc* or *Schizothrix* (Weber et al., 2016). These genera found in our study area also enhance the ecological role of biocrusts. For example, *Microcoleus* has a thick EPS layer that surrounds its trichomes in bundles and can remain over many years after thricomes die, contributing to bioestabilization of soils (Büdel et al., 2016). Cyanobacteria adapted to live within the biocrust as *Scytonema* are indicators of the state of maturity of this biocrusts, with a thick and well developed structure dominated by lichens and mosses (Büdel et al., 2016). *Scytonema* is able to enhance microhabitat conditions in exposed bare ground areas by producing a UV protector pigment called scytonemin (Sinha and Häder, 2008). Overall, heterocyst-forming cyanobacteria found such as *Mojavia*, *Roholtiella*, *Scytonema* or *Nostoc* perform N fixation and are important contributors as well to C fixation in these soils (Belnap, 2002). Our study contributes to our understanding of biocrust-associated cyanobacteria from Mediterranean areas, and describes two new phylotypes of *Microcoleus* and *Scytonema*, two of the most important cyanobacterial genera found in dryland soils worldwide.

## Acknowledgements

This work was supported by grants from the Ministerio de Economía y Competitividad, Spain (CGL-2013-44870-R and CGL2013-44661-R). FTM and CCD acknowledge support from the European Research Council (ERC Grant Agreement 647038 [BIODESERT]).

## References

Altschul, S.F., Gish, W., Miller, W., Myers, E.W., Lipman, D.J., 1990. Basic local alignment search tool. J. Mol. Biol. 215, 403–410.

Anagnostidis, K., Komárek, J., 1999. Süsswasserflora von Mitteleuropa: Cyanoprokaryota; Teil 1: Chroococcales, Süsswasserflora von Mitteleuropa. Fischer, Jena.

Belnap, J., 2002. Nitrogen fixation in biological soil crusts from southeast Utah, USA. Biol. Fertil. Soils 35, 128–135. doi:10.1007/s00374-002-0452-x

Berrendero, E., Perona, E., Mateo, P., 2011. Phenotypic variability and phylogenetic relationships of the genera *Tolypothrix* and *Calothrix* (Nostocales, Cyanobacteria) from running water. Int. J. Syst. Evol. Microbiol. 61, 3039–3051. doi:10.1099/ijs.0.027581-0

Bohunická, M., Pietrasiak, N., Johansen, J.R., Gómez, E.B., Hauer, T., Gaysina, L.A., Lukešová, A., 2015. *Roholtiella*, gen. nov. (Nostocales, Cyanobacteria)—A tapering and branching cyanobacteria of the family Nostocaceae. Phytotaxa 197, 84–103. doi:10.11646/phytotaxa.197.2.2

Bowker, M.A., Maestre, F.T., Mau, R.L., 2013. Diversity and Patch-Size Distributions of Biological Soil Crusts Regulate Dryland Ecosystem Multifunctionality. Ecosystems 16, 923–933. doi:10.1007/s10021-013-9644–5

Büdel, B., Dulić, T., Darienko, T., Rybalka, N., Friedl, T., 2016. Cyanobacteria and Algae of Biological Soil Crusts, in: Weber, B., Büdel, B., Belnap, J. (Eds.), Biological Soil Crusts: An Organizing Principle in Drylands. Springer International Publishing, Cham, pp. 55–80. doi:10.1007/978-3-319-30214-0_4

Castillo-Monroy, A.P., Maestre, F.T., Delgado-Baquerizo, M., Gallardo, A., 2010. Biological soil crusts modulate nitrogen availability in semi-arid ecosystems: Insights from a Mediterranean grassland. Plant Soil 333, 21–34. doi:10.1007/s11104-009-0276-7

Dojani, S., Kauff, F., Weber, B., Büdel, B., 2014. Genotypic and Phenotypic Diversity of Cyanobacteria in Biological Soil Crusts of the Succulent Karoo and Nama Karoo of Southern Africa. Microb. Ecol. 67, 286–301. doi:10.1007/s00248-013-0301-5

Elbert, W., Weber, B., Burrows, S., Steinkamp, J., Büdel, B., Andreae, M.O., Pöschl, U., 2012. Contribution of cryptogamic covers to the global cycles of carbon and nitrogen. Nat. Geosci. 5, 459–462. doi:10.1038/ngeo1486

Escudero, A., Palacio, S., Maestre, F.T., Luzuriaga, A.L., 2014. Plant life on gypsum: A review of its multiple facets. Biol. Rev. 90, 1–18. doi:10.1111/brv.12092

Felsenstein, J., 1985. Confidence limits on phylogenies: an approach using the bootstrap. Evolution (N. Y). 783–791.

Flechtner, V.R., Boyer, S.L., Johansen, J.R., DeNoble, M.L., 2002. *Spirirestis rafaelensis* gen. et sp. nov. (Cyanophyceae), a new cyanobacterial genus from arid soils. Nov. Hedwigia 74, 1–24. doi:10.1127/0029-5035/2002/0074-0001

Garcia-Pichel, F., López-Cortés, A., Nübel, U., 2001. Phylogenetic and Morphological Diversity of Cyanobacteria in Soil Desert Crusts from the Colorado Plateau. Appl. Environ. Microbiol. 67, 1902–1910. doi:10.1128/AEM.67.4.1902-1910.2001

Garcia-Pichel, F., Loza, V., Marusenko, Y., Mateo, P., Potrafka, R.M., 2013. Temperature Drives the Continental-Scale Distribution of Key Microbes in Topsoil Communities. Science (80-.). 340, 1574–1577. doi:10.1126/science.1236404

Garcia-Pichel, F., Wojciechowski, M.F., 2009. The evolution of a capacity to build supra-cellular ropes enabled filamentous cyanobacteria to colonize highly erodible substrates. PLoS One 4, 4–9. doi:10.1371/journal.pone.0007801

Geitler, L., 1932. Cyanophyceae, in: Cyanophyceae. Johnson, New York, p. 1196.

Gkelis, S., Rajaniemi, P., Vardaka, E., Moustaka-Gouni, M., Lanaras, T., Sivonen, K., 2005. Limnothrix redekei (Van Goor) Meffert (Cyanobacteria) strains from Lake Kastoria, Greece form a separate phylogenetic group. Microb. Ecol. 49, 176–182. doi:10.1007/s00248-003-2030-7

Gómez, N., Charles Donato, J., Giorgi, A., Guasch, H., Mateo, P., Sabater, S., 2009. La biota de los ríos: los microorganismos autótrofos. Conceptos y técnicas en Ecol. Fluv. 219–247. doi:10.1017/CBO9781107415324.004

Hagemann, M., Henneberg, M., Felde, V.J.M.N.L., Drahorad, S.L., Berkowicz, S.M., Felix-Henningsen, P., Kaplan, A., 2014. Cyanobacterial Diversity in Biological Soil Crusts along a Precipitation Gradient, Northwest Negev Desert, Israel. Microb. Ecol. 219–230. doi:10.1007/s00248-014- 0533-z

IUSS Working Group WRB, 2014. World reference base for soil resources 2014. International soil classification system for naming soils and creating legends for soil maps, World Soil Resources Reports No. 106. doi:10.1017/S0014479706394902

Komárek, J., 2013. Süsswasserflora von Mitteleuropa: Cyanoprokaryota?; Teil 3: Heterocytous genera, Süsswasserflora von Mitteleuropa. Springer, Berlin.

Komárek, J., Anagnostidis, K., 2005. Süsswasserflora von Mitteleuropa: Cyanoprokaryota; Teil 2: Oscillatoriales, Süsswasserflora von Mitteleuropa. Elsevier, München.

Kumar, D., Adhikary, S.P., 2015. Diversity, molecular phylogeny, and metabolic activity of cyanobacteria in biological soil crusts from Santiniketan (India). J. Appl. Phycol. 27, 339–349. doi:10.1007/s10811-014-0328-0

Loza, V., Perona, E., Mateo, P., 2013. Molecular fingerprinting of cyanobacteria from river biofilms as a water quality monitoring tool. Appl. Environ. Microbiol. 79, 1459–1472. doi:10.1128/AEM.03351–12

Lyra, C., Suomalainen, S., Gugger, M., Vezie, C., Sundman, P., Paulin, L., Sivonen, K., 2001. Molecular characterization of planktic cyanobacteria of Anabaena, Aphanizomenon, Microcystis and Planktothrix genera. Int. J. Syst. Evol. Microbiol. 51, 513–526. doi:10.1099/00207713-51-2-513

Maestre, F.T., Bowker, M.A., Cantón, Y., Castillo-Monroy, A.P., Cortina, J., Escolar, C., Escudero, A., Lázaro, R., Martínez, I., 2011. Ecology and functional roles of biological soil crusts in semi-arid ecosystems of Spain. J. Arid Environ. 75, 1282–1291. doi:10.1016/j.jaridenv.2010.12.008

Maestre, F.T., Escolar, C., de Guevara, M.L., Quero, J.L., Lázaro, R., Delgado-Baquerizo, M., Ochoa, V., Berdugo, M., Gozalo, B., Gallardo, A., 2013. Changes in biocrust cover drive carbon cycle responses to climate change in drylands. Glob. Chang. Biol. 19, 3835–3847. doi:10.1111/gcb.12306

Maestre, F.T., Martín, N., Díez, B., López-Poma, R., Santos, F., Luque, I., Cortina, J., 2006. Watering, fertilization, and slurry inoculation promote recovery of biological crust function in degraded soils. Microb. Ecol. 52, 365–377. doi:10.1007/s00248-006-9017-0

Maestre, F.T., Quero, J.L., Gotelli, N.J., Escudero, A., Ochoa, V., Delgado-Baquerizo, M., García-Gómez, M., Bowker, M. a, Soliveres, S., Escolar, C., García-Palacios, P., Berdugo, M., Valencia, E., Gozalo, B., Gallardo, A., Aguilera, L., Arredondo, T., Blones, J., Boeken, B., Bran, D., Conceição, A.a, Cabrera, O., 2012. Plant Species Richness and Ecosystems Multifunctionality in Global Drylands. Science (80-.). 335, 2014–2017. doi:10.1126/science.1215442

Mager, D.M., Thomas, A.D., 2011. Extracellular polysaccharides from cyanobacterial soil crusts: A review of their role in dryland soil processes. J. Arid Environ. 75, 91–97. doi:10.1016/j.jaridenv.2010.10.001

Martínez, I., Escudero, A., Maestre, F.T., De La Cruz, A., Guerrero, C., Rubio, A., 2006. Small-scale patterns of abundance of mosses and lichens forming biological soil crusts in two semi-arid gypsum environments. Aust. J. Bot. 54, 339–348. doi:10.1071/BT05078

Neilan, B.A., Jacobs, D., Therese, D.D., Blackall, L.L., Hawkins, P.R., Cox, P.T., Goodman, A.E., 1997. rRNA Sequences and Evolutionary Relationships among Toxic and Nontoxic Cyanobacteria of the Genus Microcystis. Int. J. Syst. Bacteriol. 47, 693–697. doi:10.1099/00207713-47-3-693

Perkerson III, R.B., Johansen, J.R., Kovácik, L., Brand, J., Kaštovský, J., Casamatta, D.A., 2011. A unique Pseudanabaenalean (Cyanobacteria) genus Nodosilinea gen.nov. based on morphological and molecular data. J. Phycol. 47, 1397–1412. doi:10.1111/j.1529-8817.2011.01077.x

Reháková, K., Johansen, J.R., Casamatta, D.A., Xuesong, L., Vincent, J., 2007. Morphological and molecular characterization of selected desert soil cyanobacteria: three species new to science including Mojavia pulchra gen. et sp. nov. Phycologia 46, 481–502.

Rippka, R., Deruelles, J., Waterbury, J.B., Herdman, M., Stanier, R.Y., 1979. Generic Assignments, Strain Histories and Properties of Pure Cultures of Cyanobacteria. J. Gen. Microbiol. 111, 1–61. doi:10.1099/00221287-111-1-1

Saitou, N., Nei, M., 1987. The neighbor-joining method: a new method for reconstructing phylogenetic trees. Mol. Biol. Evol. 4, 406–425.

Sinha, R.P., Häder, D.P., 2008. UV-protectants in cyanobacteria. Plant Sci. 174, 278–289. doi:10.1016/j.plantsci.2007.12.004

Steven, B., Gallegos-Graves, L.V., Belnap, J., Kuske, C.R., 2013. Dryland soil microbial communities display spatial biogeographic patterns associated with soil depth and soil parent material. FEMS Microbiol. Ecol. 86, 101–113. doi:10.1111/1574-6941.12143

Tajima, F., Nei, M., 1984. Estimation of evolutionary distance between nucleotide sequences. Mol. Biol. Evol. 1, 269–285.

Tamaru, Y., Takani, Y., Yoshida, T., Sakamoto, T., 2005. Crucial Role of Extracellular Polysaccharides in Desiccation and Freezing Tolerance in the Terrestrial Cyanobacterium *Nostoc commune*. Appl. Environ. Microbiol. 71, 7327–7333. doi:10.1128/AEM.71.11.7327

Tamura, K., Stecher, G., Peterson, D., Filipski, A., Kumar, S., 2013. MEGA6: molecular evolutionary genetics analysis version 6.0. Mol. Biol. Evol. 30, 2725–2729.

Thomazeau, S., Houdan-Fourmont, A., Coute, A., Duval, C., Couloux, A., Rousseau, F., Bernard, C., 2010. The Contribution of Sub-Saharian African Strains to the Phylogeny of Cyanobacteria: Focusing on the Nostocaceae (Nostocales, Cyanobacteria). J. Phycol. 46, 564–579.

Thompson, J.D., Higgins, D.G., Gibson, T.J., 1994. Clustal-W - Improving the Sensitivity of Progressive Multiple Sequence Alignment Through Sequence Weighting, Position-Specific Gap Penalties and Weight Matrix Choice. Nucleic Acids Res. 22, 4673–4680. doi:10.1093/nar/22.22.4673

Weber, B., Belnap, J., Büdel, B., 2016. Synthesis on Biological Soil Crust Research, in: Weber, B., Büdel, B., Belnap, J. (Eds.), Biological Soil Crusts: An Organizing Principle in Drylands. Springer International Publishing, Cham, pp. 527–534. doi:10.1007/978-3-319-30214-0_25

Williams, L., Loewen-Schneider, K., Maier, S., Büdel, B., 2016. Cyanobacterial diversity of western European biological soil crusts along a latitudinal gradient. FEMS Microbiol. Ecol. 92, fiw157. doi:10.1093/femsec/fiw157

Yeager, C.M., Kuske, C.R., Carney, T.D., Johnson, S.L., Ticknor, L.O., Belnap, J., 2012. Response of biological soil crust diazotrophs to season, altered summer precipitation, and year-round increased temperature in an arid grassland of the Colorado Plateau, USA. Front. Microbiol. 3. doi:10.3389/fmicb.2012.00358

